# Bioinformatics Study of Genetic Variability in Obelisks

**DOI:** 10.1101/2025.02.24.639975

**Authors:** Francisco Javier Hernández-Walias, Roberto Reinosa-Fernández

## Abstract

**Motivation:** The importance of in silico genetic variability analyses of new biological entities, such as obelisks, lies in their potential to serve as a starting point for experimental studies. The study of obelisks could open new horizons in the field of biological sciences.

**Results:** Several findings have been determined, including high conservation among the sequences within each cluster, despite the presence of significant nucleotide mutations; high variability among the Oblin-1 proteins found in the consensus sequences of each cluster, despite all sharing a known conserved region; structural consistency in the 3D model of the Oblin-2 consensus with previous studies; a series of nucleotide and protein-level consensus sequences for each cluster, where nucleotide-level patterns show color-coded conservation percentages that could be used in further research, such as primer or probe design, and protein-level sequences are presented in FASTA format for Oblin- 1, Oblin-2, and unknown Open Reading Frames (ORFs).

## Introduction

### Nucleotide pattern

Certain biological entities have been proposed by Zheludev et al., 2024, under the name “obelisks,” as potential new discoveries in the microbiome, including the human microbiome. Approximately 30,000 obelisks have been identified based on their homology in ribonucleotide sequences. The known obelisks are around 1,000 nucleotides in size and exhibit various ribozyme-related variants, particularly the Type III self-cleaving ribozyme (Zheludev et al., 2024).

This group has analyzed the nucleotide-level homology between obelisks and known genomes or viromes, including viroids and virusoids, and found low sequence similarity. However, they may share certain structural and/or functional similarities. Obelisks have a circular genomic structure that folds upon itself, forming a distinct elongated spiral conformation. At the protein level, Zheludev et al., 2024, determined that obelisk RNA sequences encode an exclusive group of proteins termed “Oblin.” No homologous structural folds have been identified between these proteins and those derived from previously known messenger RNAs in genomes and viromes, but bioinformatics simulations of their folding have been conducted.

Aside from virome analyses, virology studies viroid and virusoid gene sequences, which show low similarity to obelisks. Some virome sequences function as internal promoters known as Internal Ribosome Entry Sites (IRES), which are RNA regions that facilitate translation. IRES elements remain functional under conditions where Cap-dependent RNA translation is blocked (Lee et al., 2017). In the case of positive-sense single-stranded RNA viruses, their own sequences act as messenger RNAs for IRES-mediated translation, serving as ribosomal recognition sites and forming secondary structures morphologically similar to viroids (Yang et al., 2019).

Due to the rod-shaped secondary structure of obelisks, studying their conformation in various structural domains of IRES elements could provide insights into the translational or even transcriptional capacity of viroids, obelisks, and possibly some virusoids. This intriguing similarity between the secondary structures of IRES elements and the gene sequences of these biological entities may help to explain the presence of similar nucleotide loops in all of them.

The ribozyme-based replication machinery of obelisks, however, appears to resemble viroids, particularly those with Type III or “hammerhead” ribozyme sequences. The hammerhead ribozyme structure has been studied as a replicative sequence (Flores et al., 2000; Giguère et al., 2014). As an example of a ribozyme present in the studied obelisks, ObV-HHR3 was identified, resembling viroids from the Avsunviroidae family. Furthermore, a functional relationship between this ribozyme and Oblin-1 was proposed, as these ribozyme sequences encode divergent Oblin-1 proteins but not Oblin-2 (Zheludev et al., 2024). Additionally, the functional involvement of Oblin-1 and Oblin-2 was established as a key part of obelisk replication.

The difficulty in assigning a definitive similarity between obelisks and virusoids, such as the hepatitis delta virus, stems from the latter’s infectious collaboration as a satellite entity of a virus. The viroid itself acts as an “auxiliary virus” by incorporating surface antigens from its reference virus, which, in the case of the hepatitis delta virus, is the hepatitis B virus (Chiou et al., 2022). In contrast, no antigenic protein associations have been found in obelisks.

### Protein pattern

Two known obelisks, designated alpha (1164 nt) and beta (1182 nt), are capable of encoding at least two types of oblins: Oblin-1 and Oblin-2. Specifically, in Obelisk-alpha, Oblin-1 is encoded as a 202-amino-acid protein, while Oblin-2 is encoded as a 53-amino-acid protein.

#### Oblin-1

The structural conformation of Oblin-1 remains unknown, but the estimation by Zheludev et al., 2024, suggested a globular structure with alpha-helices and beta-sheets.

#### Oblin-2

Findings by Zheludev et al., 2024, estimate a conformational distribution with an alpha-helix forming a leucine zipper motif.

### Bacterial Population as a Host for Obelisks

In bacterial populations of Streptococcus sanguinis (strain SK36) from the human digestive microbiome, obelisk RNA sequences have been identified, though their physiological impact on the microbiome or the host organism remains unknown (Schmitt- Kremer et al., 2024).

A specific obelisk sequence, “Obelisk S.s,” was found in this strain, measuring 1,137 nucleotides in length, with genomic similarity to alpha and beta obelisks of 41% and 35%, respectively. No genomic similarities to Oblin-2 have been found, although similarities to Oblin-1 exist.

The presence of obelisk sequences in bacterial populations, at least in the mentioned strain, suggests that they may exist in other strains or even species. It remains unclear whether different types of obelisks exhibit tropism for eukaryotic cells (though they seem ecologically related to microbial populations and their host organisms (Schmitt-Kremer et al., 2024)), whether they provide a beneficial function to bacteria (Maddamsetti et al., 2024), or whether there are facilitated mechanisms of obelisk sequence transmission to the cited strain. It has been proposed that bacterial transport via vesicles containing small RNAs might serve as a communication strategy that could be key to a potential mechanism of obelisk transmission (Maddamsetti et al., 2024).

However, the presence of obelisks in cellular hosts seems relevant for protein production, specifically for at least two types (alpha and beta obelisks). As potential new biological entities, it remains unknown whether they possess a translational mechanism similar to bacteriophages or an autoreplicative mechanism akin to viroids, pending further studies on whether the protein structures predicted by bioinformatics tools are actually generated.

Based on the hypotheses proposed by Zheludev et al., 2024, it could be speculated that, as with cellular genomic evolution incorporating external sequences into current cellular sequences, the obelisk S.s. might contribute to transcriptomic fragments in Streptococcus sanguinis, as suggested by bioinformatics analyses detecting obelisk sequences in bacterial transcriptomes (Schmitt-Kremer et al., 2024), potentially shaping the current transcriptomic sequence of strain SK36.

## Methods

The sequences of the obelisks were initially obtained from the work of Zheludev et al., 2024, available in Stanford’s digital repository (https://purl.stanford.edu/wb363nt3637). These sequences were stored in a CSV file, where each row contained relevant information for each obelisk. Using this data, a FASTA file was generated, where the header included the corresponding ID and cluster, followed by the sequence on the next line. This process was carried out using a small Python script (Van Rossum et al., 2009). Next, the sequences were separated by cluster, yielding 26 files. One of these corresponded to sequences whose cluster was indicated as “.” and was therefore discarded. This entire process was performed using the header filtering function of the EpiMolBio software (Reinosa et al., 2025).

Subsequently, a multiple sequence alignment was conducted using the corresponding function of EpiMolBio (Reinosa et al., 2025), which employs MUSCLE (Edgar, 2004) for the alignment. The resulting files were opened in MEGAX (Kumar et al., 2018) to remove insertions, discarding those present in more than 50% of the sequences.

With the curated files, consensus sequences were calculated using the conservation function of EpiMolBio (Reinosa et al., 2025). Due to the low number of sequences in some clusters, some consensus sequences were either non-functional or merely replicated the only representative sequence; thus, clusters with fewer than 10 sequences were discarded, reducing the total from 25 to 12.

Once the consensus sequences were obtained, they were used as references to identify mutations present in each cluster using the mutation table in the polymorphism section of the EpiMolBio software (Reinosa et al., 2025). Additionally, conservation and mutation frequency were calculated using the corresponding functions of this software.

At this point, a bioinformatics software developed by Reinosa, 2017, originally designed for ORF detection in retroviruses, was employed. This software is suitable for identifying ORFs in biological entities with a standard genetic code and small genomes. The consensus sequences of each cluster were input into the program to attempt the identification of Oblin-1 and Oblin-2.

Candidates compatible with the Oblin-1 protein, obtained from the consensus sequences of each cluster, were aligned using MUSCLE (Edgar, 2004) included in MEGAX (Kumar et al., 2018) to identify conserved regions. Finally, the 3D structures of the Oblin-2 protein from the alpha cluster consensus were predicted using AlphaFold3 (Abramson et al., 2024) for result presentation.

## Results

### Mutations

This section presents mutations with a frequency above 20%, the total number of sequences in each cluster, and the overall number of sequences. Insertions and deletions were not considered in this analysis:

**Table 1.**
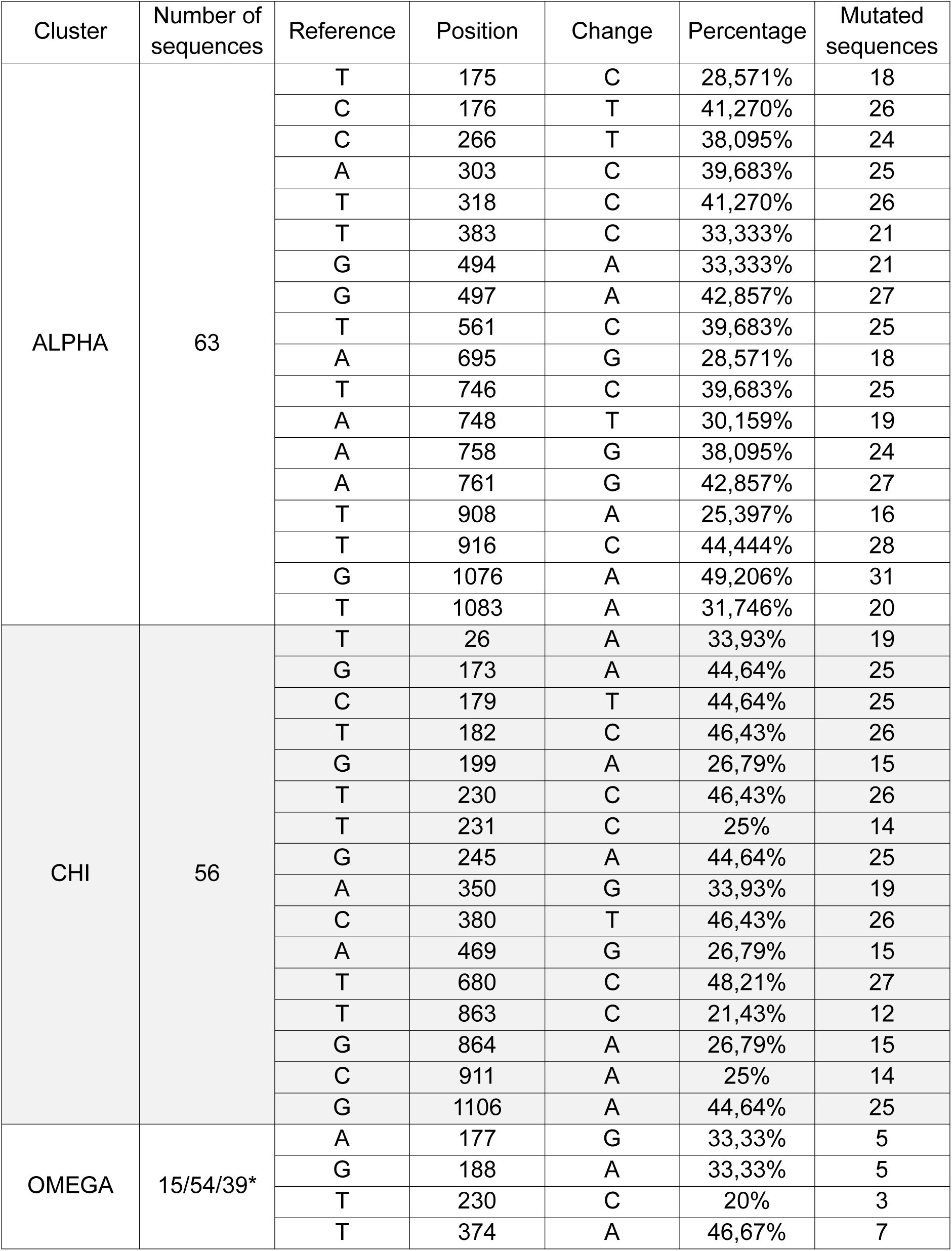
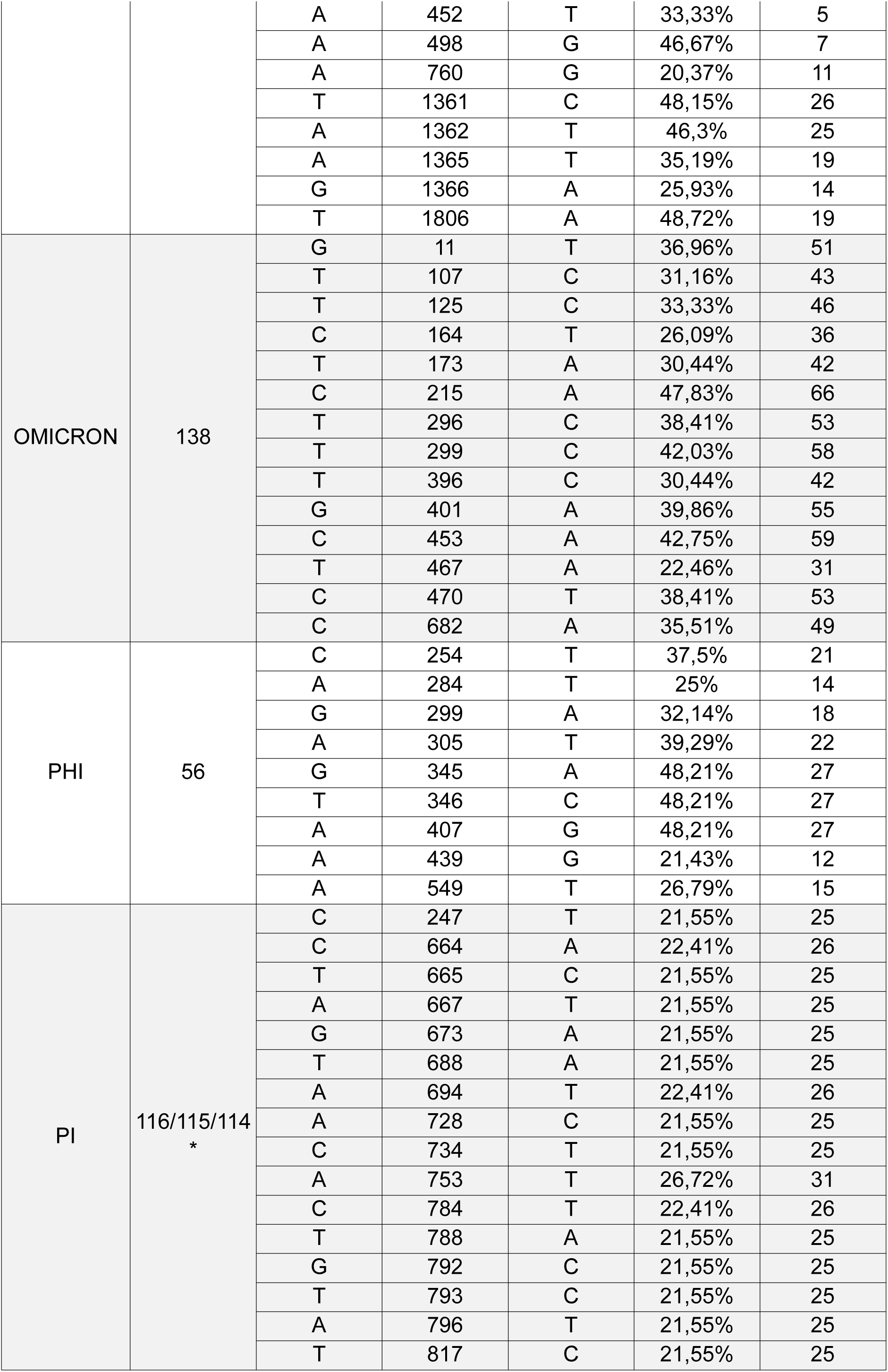
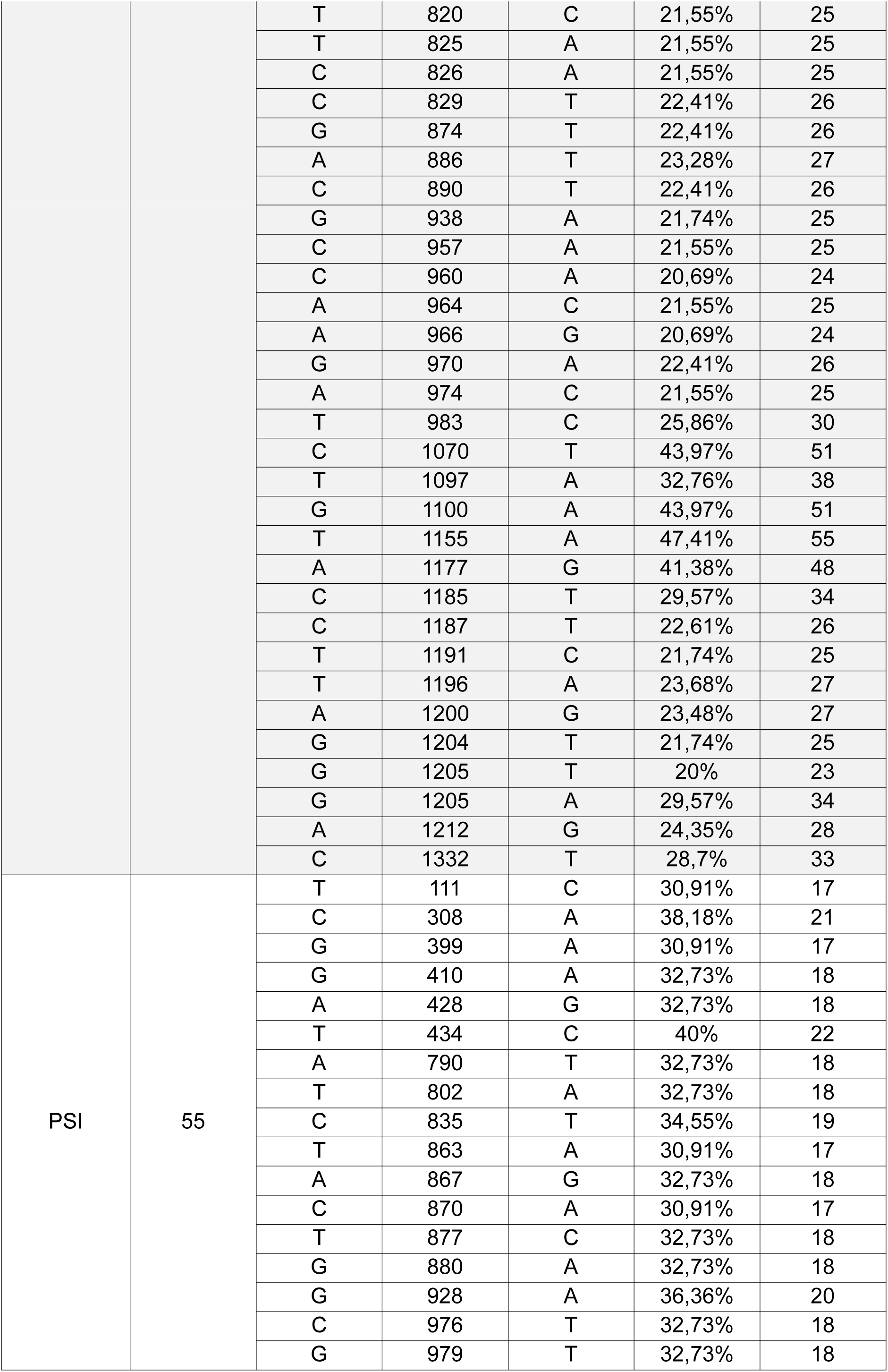
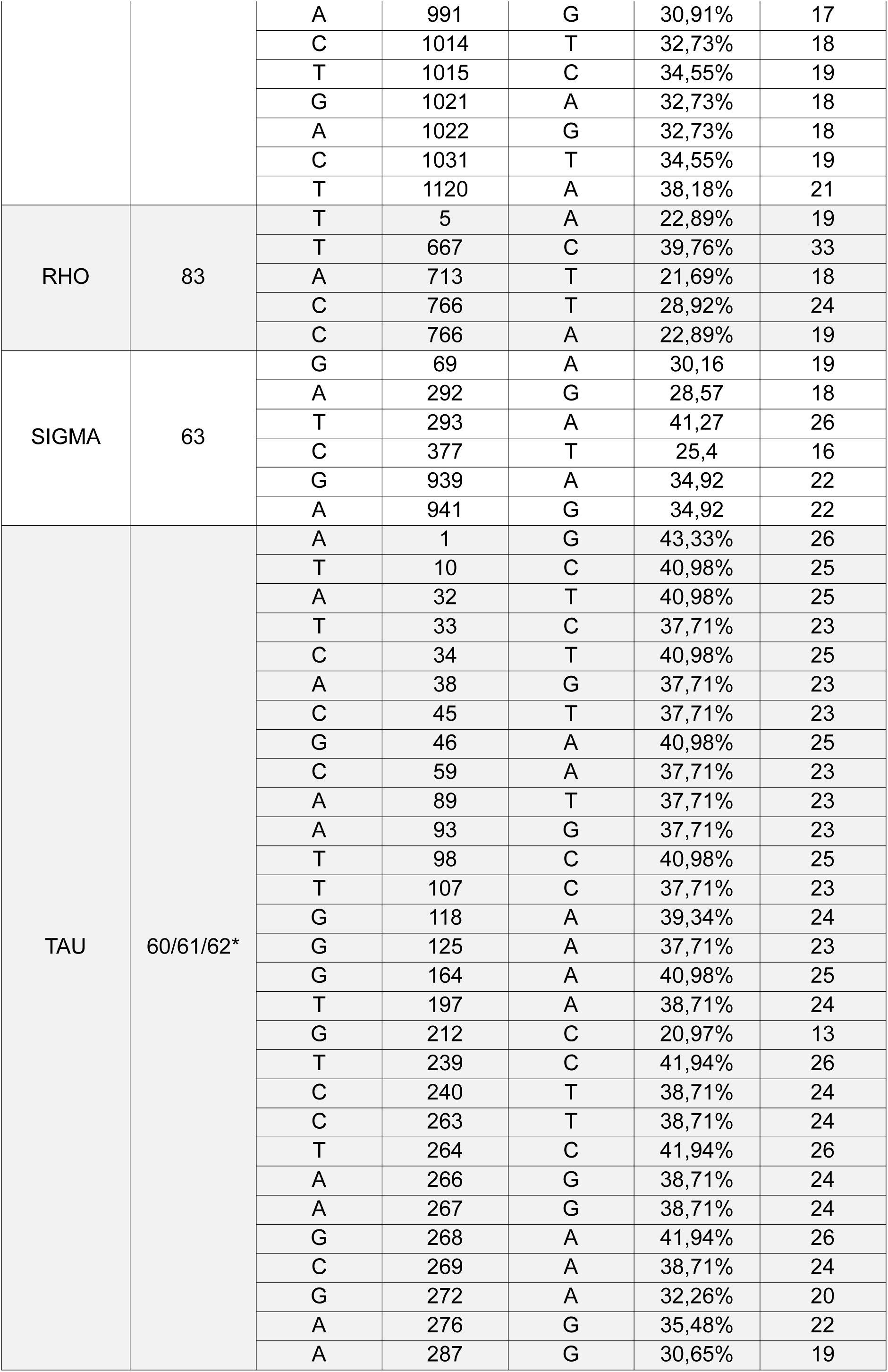
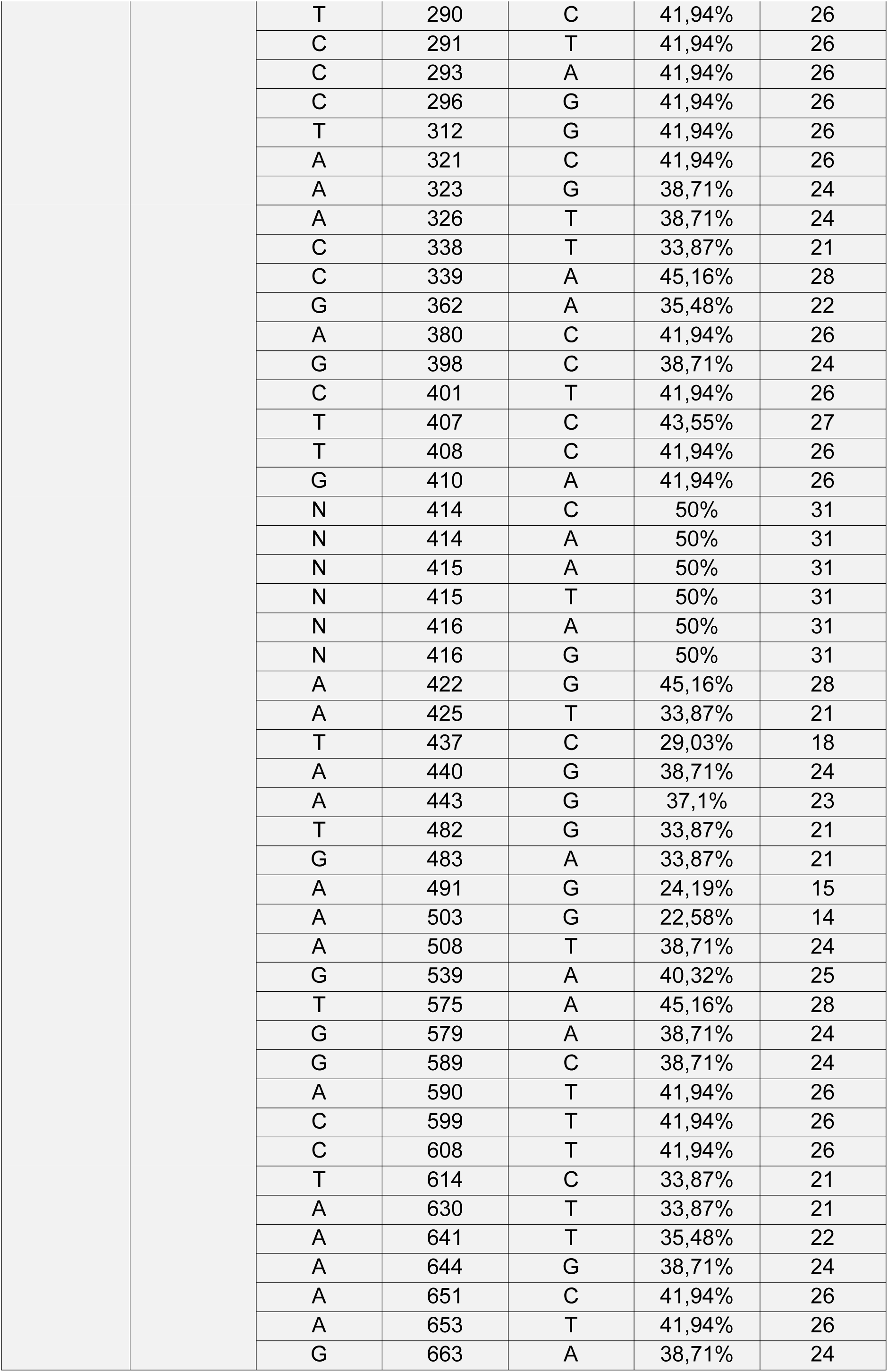
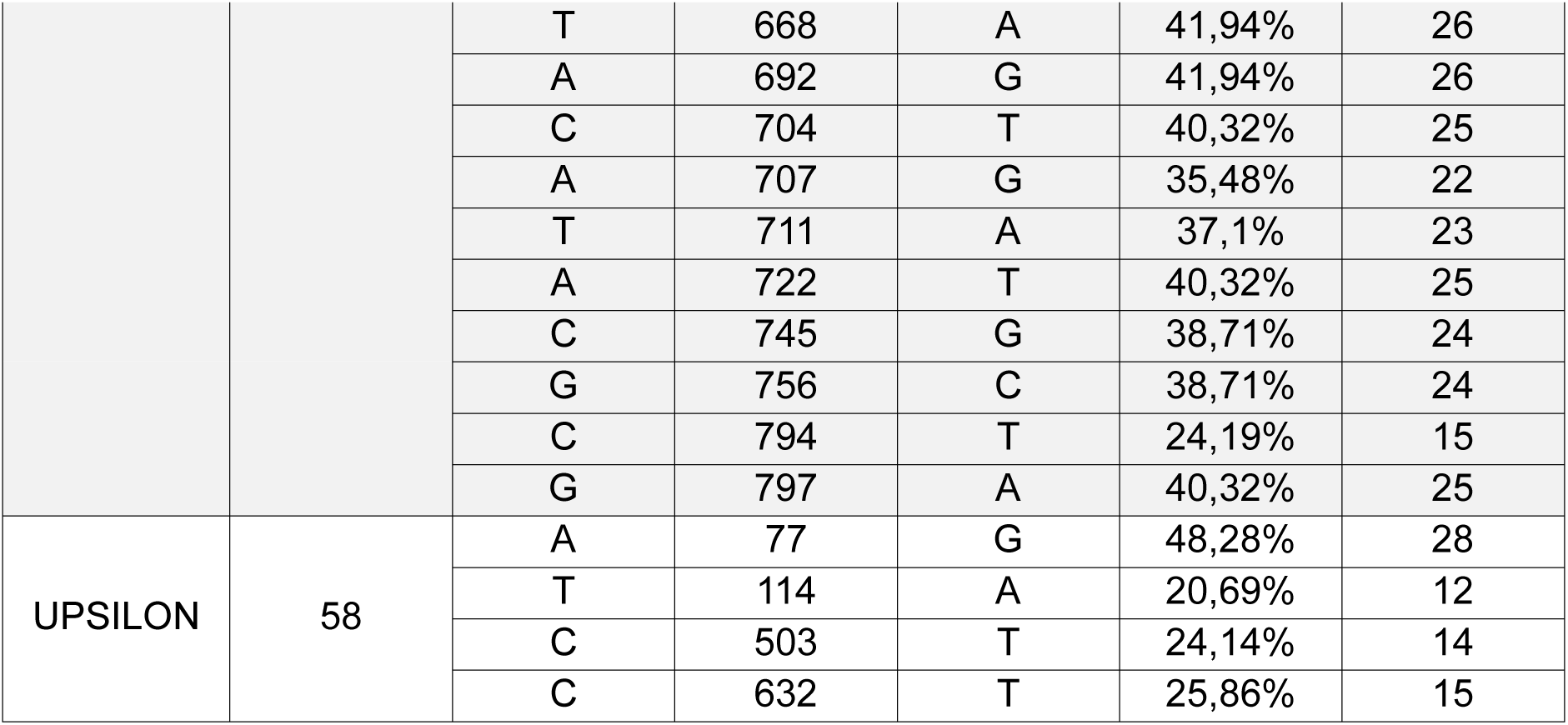
Mutation analysis of each cluster. (*Cluster with a different number of sequences at each position. This may be due to the presence of deletions and/or incomplete sequences).

### Mutation frequency

This section presents a table displaying the mutation frequency for each cluster, the conservation percentage, and the average number of mutated sequences per cluster:

### Consensus Sequences

Consensus sequences generated for each cluster. The color code corresponds to a conservation pattern as follows:

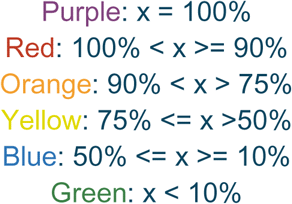

#### ALPHA

TCACTTTCTGAAACCTATCTACCAACAACTTAATAGAAAGGAACAGTGAAGTGAGAGACATTGAACTTGATTCCAGTGCTTTTAGGAGTCAAGTTTCTCTACTGAGTCAAGAAACTTCAGAGAAATTTCTCACAGGAGCTGCCCTTGTAAGTCCCAAGCGTTCGAAATACTACATCTCGGAGGTAGAAGGGCTCAAAGTGCACAGCAGATCGAAGAAAGACCTTCTTGCTCTTGCTATCATAAGTTGGTGGTTAGAAGATTCCATCAGATTCTATCTGCAAGAGGAATTGTACTTTCTAAGTATTAACAATTCTGATTTAATTGAAATTAGACTTTGTCTAACTTCAAAGTCAGGAATGTTAAATTTCTTAGAAGATACAACTCTCTATCACAGTAGAGATCTTTTTGGAAATATTCTTCCTACCAGCCCTGAAAAGCAAGTAAGATTGGCCAACCTCGTCAGCGTACGCTATGGACCAACTTCTTTACCGAAGAGGGTAATTCGACGTCGTGGCTACAAGGACCATGGCTCGAGGAGATTTCCTCATGAAGTTCATGACTTAAGTAGTGGAAAACTTGCCCAAATTAAGTACGAAGAAGAAATTCAATCCTATCACGATACTCTCCTTTTCTTAAGAGGCTGGTTAGACGGATTCTAGATTTGGAACTCCGGAACTCGCTCTAATTAGTTTTCACTGAAAGGAGACAAAGTTATGGATTCTGTTCAAATTCTAAGGAAGAAGATTCAAAAGAATGAAGAACAAAGAGAGTTTCTTCTAAAGAAGATTGGGAATCTTGAATATGAGATTAACAATCTTGAACACAAAATTGAAAATCAGCAAAGAGTGCTTCAGAACCTACTAAAGGAGAAGTAGAGCGTTGCCGGTTAGGTTAATCCCCGAAGTTATTAGACCTTTCGGTCGAAAGCTTACGGGGATTACCCCGATATAGCAATCCTCTGCTTTCTGACTAAGTAAGGTCAGCTGCAGACTTGAGAGTTGACAATCTTGTATTCAACTGTCAATCGCTATATTAAAGATACCTTTGATTCTAAAGAGGAATCTCTCTGTTCCTCGATCAAATGTGTCTTCAACCTGGAAAGTGAAACAGGACCACTCCTCTGTTCCTAATCAGGAAACAGTTGAGCAATCAAGAGTTCC GGG

#### BETA

TCATTCTTCTCAAGGCTCATCCGATCATCTAAATAAGGAGACAATCAAAGATGCTAGAAGTTCGGTTTTCAACGTTAGAAAACCAGATTAGAAACTTCTATAACCTCTTTCTGAAGAAAGAGTGCGAAGATCGCAGTAGATCTGTCGTTGATAGCTCCAATCTCGAGACTGATCAGTTTGAACCGAACGGTTTATCTTTTGCTTCTAACTTAAGTCTAACAGAGAAAATTCTTCTCGGAGTCTTCCAATGGTATATGCCGAAAGAACTCGGTGTTCTTACGAATCTTTGGTTAGAGGAACACTGGGGCGGTGAATTTAAAGAAATAAAAGCAGTGCTTTTAAATTCTAAAGACACCGCTCTAGGATATCTCCTAGTCTCGGATCGTTGGTCCACCAGAGATTTTTACGGAAATATTCTAAAGGAAAATAACGTGAAGAGGATTCTCCTGTCGATGAAAGTTAGACGTAGAAATAAATCTAGGCCCAAACGAAAGGTCT GGAGACGTGGCTACAACGACAAAGGATCCACACGTCTTCCGCACGATTCTGTCAGATTCGATTATAGAAAATTAAAATCTGTTATTCAGACGATTGAAGAACAGGAACTCCTGGACTCTAAGATAGAGTCCTTATTTTTCATAATCGGACAAAGAATTGAGAAAGACGAAATCCTGGACTACTACGACTCAATAGAGTTCTAATTAGGTTCCTAGGCGATAGCGGGGTAGGATTTTCCACCCCACTATCTTTTGTTCTTTTTAAAGAAAGAAAGAGAAAATTTAATTAGAGAGGTATAAGATTATGACGAAACGTGAAACGAAGATCCAGAATCTTCAGAAGCTAGTCGAAGACATCGAGAAGAAAGTGGAGAAACTAGAGGCTCAGAAGAAAGTCTATTTACTGACAATCGAAAGTCTCAAATCAAAACTTGATTCGAAATCTGACGATGAAAGAAAATAGAGTTCTAGAAAGCTATCTAGAATTCCTCGATCTTCTGACTCTACAACTAGAGTCAGGAGATCTGTTTCTTACATTTGACGATTCGAACATGATGATATACCACTTAAGAAAATTCTCTCGACTTCTTTTAAAAGCACAGGATAGTGTTCGGAGTGGAAATCACAGTCCTCTATTGTCAAAGAAGACAAATCAGATCTATATTAAGTTGTATGATCTAGGACT

#### CHI

TAAGTATTAGTTGTTGTTCTCGCAATTCAACTAAGAAAGGATGATTCAGAATGCGAGATAAGAATCTAACTCTTAAACAATTTAATATTCAGTTGGAGAGAAAAATTCATCAAGGTAATCCCTTTGATGAAAGGTTTCTCACTAGACCTGTCTTTGTAAGTCCGACACGACCGATTGACCCTTCAGAGGTTTCAATACGATATCCTCGAGATTTTAGACAGTTGAGAAGTTTGTGTATCTTACAGTGGTATGTACCAGAAACACTACACTGGAGAATGTTCCTAGATCTAAAGGACTTTTCATTTTCTCAGTTGAACGAAAAGCAAAGGATAGAAATCCAAATACTTTTAAGTTCAAAAGAAGATATGATCAGTTATCTCTATTTCACAGAACGATACTCAGGAAGTGAAATTTTTGGAAACATACTCGGTAATGATCTCACAGATCTCAAAAAGTCAATGATCTTCAAAAGATTGAAGTATAAAAGACCAAAGAGAGCAATTCGTCGCCGTGGCTACAAAGACAAAGGGTCCAGGAGACCAGATCATCGGTGGCTACCAAAAGAAGATTTCACCTTCACCGAATTGCAAATTGAAAAAGAAAAAAGAAACTATCAACTACAAACTATCACAAACCGTATCTTGAAGACTTTGAGGCAACAGCTAAAAAGCTAAAGATTTTCGGAGATAGCGAGTACTGGTAACTTCGTTCTCAATTTCCTTAGATTGAATTTATCGCGACGTCTCAGTTAAGATCCAGGATCCACATCCTAGTATCAAGAGGAGGTGAAGAAATCAATGCGAGTCGATCTACTTATTCGGTATCTTTCTACGAGTATAAATCGATTACAATTAGTTTCAAATGTTCTAATGAAAGAAAATCAGAAGCTAACAGCAAGAAACTCAAACAACGAGAATCTTGACGATACCCGGCACGACCAAATAACAGGAATACTAAAAGAATATAACGGCGATGCCATTAATCCGATAGATTTACCTGTCACTGAAATCGACCTGGAAGCAGTCCGATTACTAACAGAAGAATTAGAAACAAAAGTAAGAACGATTCTTTTACTAGAAGAAAATCTTACCAAACTAATAGAAAAGGACAAGTAAAGAAATCAACACTCTCAAGATCTAGGGGTCGGAACCTCTGGATCTCTGAGATTTGTCCGAATAATTCTTTCTTCGGTAGAGAACTTGATTACTTTTCTCGGTCTCTGATCGTC

#### OMEGA

GGAGATTACTGGCTACGTCCTGAATTGTGCAAAGAAAGGAGTCCACAGATATGAAGTTGTCTTGCGATTCTGCTCTGCCCGAAACAAGAAATCTGCAAAAAGAAATTGTCAGAAATCTCACCGAGCAGTTCAAGGACCGCAGACTTCGAGGCCTTGTATTCCGACCCTCCTTGGGAAATTTTGATCCGGAACATGTAAAGTATTCGAAAAGATACCTAGATTTAGTTACTCTTCTATGTATTTGTTACTTTAGCGATCTCCTGGACCCATTGGACTGGTATCTTAGAATTTCAGTTTCAGAGTACCTTTTCAAGAAACTCGATTTTCCAGA ATTGTACGCAAGTACTATTTCTAGAAGAGTTCTTTTGGAAGTTATCTCAGGAACTATTTCTACGATATCGCCGAATGTGTTCTTTGGAAACTGCTTAAGGCAACAGAATCTAGAGAGAGTACTTAAAAATTTGTATCTCTCGAGAAACTTACCTGTTAGGCTCAAAATACCCGTTAGGAGACGTGGATACAAGGACCATGGAAGTTTACGTCCCAATGAACGCTGGATTGAGAATCATGATTATTCTTTCTCAGAACTTCAGTTCGAGTTGGAACAGGAACGGCAAGATTACATCAATCTTGTGACTTCTGTTCTCAACACGATCCAGGGGTCGAGCAACAGCAATTCCTCTATTTTTGAAAATTTAAGAAAGGAGTGTCTAGAATGGCATTTGAAGAAACAAGAGATTACCTCAACATCTCAGAAAAAGTTGAGAAAAACAAACAACTGTTAGCAAAACTTGATCAGAAGATTCTTAATCTTCAGAAACAGCGAAGTGCACTTGCTGCGAAGATCACAAACCAGGAAGGTTATCTGAACAAGTGTGGAACAGGAAAGAAAATCACCAAGGCCCTTGAGATTCTCAATCTTAAGAATAAGGTCAAAGGAATTCACAAATCCTCTTCTGGAAAATCTAAGGAACTTTCTCGATAAGTAAGAGAAAATCTACACAGGGTATTCGGTTCCAGGCAGAACATTTAGAAAGAAATTTCTTATGTTGAAGAAATTCACGATCTAAGATCGTCTGTCTGGAGCCATTACTCCTGTAGCAGAATTTTCTCATTTACCAATCGAATAGTTTCAATAGTTTTCACAGAAAGGAAGGTGTGAATACCTCTAGTACCACTCTTAGGTAAAGAGAAATTCTTTAGGAACCTTGGCTACCTTCTTGTCCTATTCTACAAGATGTTCAAGAAACTTTTGTTCCTAGAAAGTTAATCTCGTAGACAATTACACATTGAACGCATTTCAGTTGATTAGAACCACATCTGACTAGTCTTAGCTCAGAAGGTCTGTCTTTCTTTCTTTTACTAGGAAGAAAAGGTAGTCTTCTAGTCTTAAAATGACATTAGATGATCTCTTTCTCTATTTTCATAGATTGGAGATTACTGGCTACGTCCTGAATTGTGCAAAGAAAGGAGTCCACAGATATGAAGTTGTCTTGCGATTCTGCTCTGCCCGAAACAAGAAATCTGCAAAAAGAAATTGTCAGAAATCTCACCGAGCAGTTCAAGGACCGCAGACTTCGAGGCCTTGTATTCCGACCCTCCTTGGGAAATTTTGATCCAGAACATGTAAAGTATTCGAAAAGATACCTAGATTTAGTTACTCTTCTATGTATTTGTTACTTTAGCGATCTCCTGGACCCATTGGACTGGTATCTTAGAATTTCAGTTTCAGAGTACCTTTTCAAGAAACTCGATTTTCCAGAATTGTACGCAAGTACTATTTCTAGAAGAGTTCTTTTGGAAGTTATCTCAGGAACTATTTCTACGATATCGCCGAATGTGTTCTTTGGAAACTGCTTAAGGCAACAGAATCTAGAGAGAGTACTTAAAAATTTGTATCTCTCGAGAAACTTACCTGTTAGGCTCAAAGTACCCGTTAGGAGACGTGGATACAAGGACCATGGAAGTTTACGTCCCAATGAACGCTGGATTGAGA ATCA

#### OMICRON

TCTTAGGGTGGCATGATCTTTGAGATTAAAAGAAAGGATGTGAACTAAAAATGCGTATCAAACTAAATAATCTAGATGTCCTATTAAATACTTCGAACTTACCTTATCGTCGTAATATCACTAGTACCGAAAGGAAACAGATCGATCGGATCTTAAATGTCTACAAGGAAGCTTATAAATACCAAGGAACCAGAATATCCGAAGTTACCTTTGACCCGAACTTTTCAGAACGAGATATTTTCATACTACTATCTTCCTTTTGTCCAGAAGTCATCGAGTTCGGTCTTCAAATTCTTTGTTTTCAGGGAGAACTAAGTCTCAACTCAGAATCAGATACAAGTATTCTATTAGGAATACTTGCCTCAGAACCGAGATTGTTACCAATTCTCCTCGAATACAAGGGTTGGAGACAGATGCCCGATGTCTCTCAGACTCTAGAAAAAATAAGAGTACAAAAGATATATCGTCCCAGAATCGAACAGGTCCAAAGAAAACGAGGATATACGGACCATGGATCTATGGCCCCTCTAGAAACTAAAATCAGAAAGAGTTGTCTCACTGACGAGTTACTAGAAGAATATCAACGACGAGTAAAATATGATCGAGTTTTAAAAGACACGATAGACTTTCTAGTCGGATACTATTCTTAGTCCACCCTCTCTCTTTAATCGAAGGTCTTCGCCACTCCTGAAAGTACACGGAGGCGATCCTCTTTAAAGAAATTATTTGAACAAAACGAAGTGGCACCCTAAGAGATACAAAGGAAATCAACATCT TTGTAGACTTTCAGGAGTGCCGAAACTTCGCCGTCAAATATCTCAATGAAGAGTAGATCCCACCGTGGC

#### PHI

CCATTGACGGAATCTCGTAGGCCAACTTAATAAAGAAAGGAGGCACTCAAGTGTCGAAAGAAGATTATTCTCTTCGACTTCTAACTTACGAAAGAGAGTTGGAACAGCTAAGCTATCCATCCTCACTTTTCGAAGATAGCCTAAAACCAGGCTCTATAGCTCCAAACTTCACTGGACAGTTTCGAAAATTCGAAGGATTTCGGGATATTGACTATGTCTATCTTGGAATGATAAGCTGGTTCTCCGATGATATCTCTCGTTGTTACTTACAGTTGTGGTTATCAGAACAGAAATCACTGTCAGGAGATTATAAAGTTCTTTTGGACCTTGTCCTTAAGAGCAAAGTAATCGCCCTGATGTGGTTTTCTGAACGATATCACACCCGTGAGTTCTTCGGAAATATCAAAAAGAGAATGCAACAAATCATCCAGATGGTCAAGCCAATAACCAAAATTCCAAGAAAACCGAAACGGTCCGTGTGGAGAAGAGGCTATAGAGACCATGGCTCCAGGCGTCCCCCCGAGAAGTGGGTGGAAACCTCAGACTGGACCTTTTCTCAATTGCAAGAAGAAATTGAGAGATATCGACAATCGCTACTTGATACTGCTGACTTTCTTGAAGGTTGGCTCTATTGATTTTCTCGCCACTGGGGGAGAAGTCGTTCTCATTCAACCATGAGTGAGAATTCTAAGTAGCCGAAGGTAATGTGGAGGAGGAACACTTCAAGTTAAAGACTAGTCTTTAACGAGTTTGGGTTCCTTCTCCCTTTACTGGGGCTTACTAGATTTCACGTCTGGAGAAGTGAAAAGATTTCTCACTAACT

#### PI

AGTATTGCTTTTCTTGTTACCAAGAAAGGGAATCAACCTCTCGGATTACTTTGCCGGAATTATTCTTCTCTGGTTCTATTGTAAGGATCTCCAGTTTACTGAGTAAAACAAACTATGGGTCTAAAACTCGGATAATTTCCCAGTACTTGGACTTTCTCAAACGTGTCAGACAAACTAAGATCCACAATAGAGTTTTATTTCTATTTATTCAACATGCGAGGTTAGCCGACATGTCATGAAAATGAACCGTCATATAGTTTCTAGTCAATTTGATTTGATGAACGTTCCAGGAGTTGAAAGAATAAAATCTATCGCCCTAGAATCCTATCAAGAAGTTGATCCAAATAAAACATTGATATCCGAGGCAGACCTTTTCGGTACAACGCTTCCCGACCTAGTGTATAAGGGGACTGTTTTAAAAAGCATCCGCCTAAGGTACACAAAGAAAGTGAAATATATCACTCTAGTCAGATTTCTTACTGGAAAAATATGTTTCGATTCAGAAAGAGTTCAGAGAATAGATCTTCTTCTCATATATGACTCTATGTTAATCCTCCAAGACTTAGTAGAGAAGGACGAAAACTTTAAAGTTAAATTTGGTTCGGATTTAGAATCCTTAGCAAAAATCCTGAAAAGTTTTAAGCTCCATCCCAAAACCAAGATCTTAGATGTGAAAAAACTAGGAAATCAAATAGAGAAAGAAATTCCTAACTTTGTCCTTCCGAAGAGAAATCTATCAACCATCTGGAAATATGTAGAGAAAATGTACTTCTTTTCTCCAAGCACCTCAAGTGGAGTCGAATTAAAACGTTTGCCTCCTAAACTCTACATTGGTAAAGGTTATACCGATAAAGGAACTGCCAGGAACCCTGCGTTAGATGGCTCACCTCCATGGCAGGAAGTAGGACAGACTTTAGCCTTTGATCCCGAGGAACCTGTAGATGAAACCAAACGAGCTACAAAAAAATAGACGATTGATTTTTAAACCTGAAGGTTACACCGCTGCTGAACTAAGAGATTGCTCCCTAGCGGATCTTCTTTTCTGTGACATGTTGAGAGAGCTCCAAAGCCACTTGGAACCTTCCAAGTTTAGACCTAAGAGTTCAAATACAAGTAAACAAAAAGAATCAAAACTCACACCAGGGTAGAGATTCTCGGTAGGTAGTCTTTGAGAGGAAGTATTGCTCTTCTTATTTCTAAGAAGGGGAATCAACCTCTCGGATTACTTTGCCGGAATTATTCTTCTCTGGTTCTATTGTAAGGATCTCCTGTTTACTGAGTAAAACAAACTTTGGGTCCAAAACTCGAATATTTTCCAGTACTTGGATCTTCCTCAAACGTGTCAGACAAACTAAGATCCAC

#### PSI

TTAGGAAGTCAAGCCAGCAACCCATTATTTCATCCCGAAAGGTGAATCTAATGAAGGAAACTGAAAAGTTACCTTCCGAAGAAACACCACTTTTGGACGACTATTTGGGTTTAGGCAAAACTGTACCTGTAACCTCATTGGAAAGAAACAATGATGCAGAGGCACAGATAGAGCCCCACCGTTTCGCCCCGAAATCCTCGAGGAGAAATGACCATACTCAGATGACGAAGACAAGTTTGCTTTACGTACTATTCGAGTATTGTATTGCGAATTCTCTCCTTGAATTCAGATTCTTTGATAGCTGTTCCACCATTTGGAAGTTGGACAAATTTGCAAATGCGAGTTTTCACTCGTCTTTCCAAAAGTCCGTTTCCGAGTGGAGAACAAAAATCATTGAAGATGAAATTTTGAGGAAAAGACTTCTCAAACAACATCGAATGGAACTAATAAGGCTTACTTATCCTTTGCCAAAAAAGCGTGGAAATTCTCCGAAGAGAAAACGCGGCTACAGTGATAAGGGCTCTACCCGACCAACGCACCAGCGCTTCCGTACCGATAAGGATACTACAGTTTCAGTTTATGCTGAAGACCTAAGAGTAGTCCGTAAAGTGAAGATCTACGGAAGGCGACCAACGGTCACCTACCAAAGACTTCCATATTCGGAAGAAATAGGTAGGTTACTGGAACTTGGTCTTCTAAGGCGAGAAGGGGACTTTATGGTCCCTATCAGACCAAAGGAAGATTAAAGGTCACGGGGTGGAACCCTTATTTCTCACACGCGAAAGGAGTAGAATTGGAAGATCCTATCCTTCAGGATCTTACCCTAGAAGAGTTCATAGAGATCTCAGACAAACTTGATCTCTCAGAAACGAAGTATCTGATTAGTATCCTAATTGTGGATCCTAATGATGATAAAGATAACTCCTCGCGGAGTTACTTCACTATCTTAGAACCACAAGAAGTCTACGATCAGGCCTCGTTCTGGAGACAAATTTGGATGGACTTCTATTTACCTTAAAGGATAAAGTCCCTCGGGAGAAGCCCTTCTTTTTCGTCCAATGGCGAAAGTGAGAAGGGTTTCACCTTTGTTGACTTTAACCTCCTTTGATTGAAGGGTACTTTGTACCCTTCAAAC

#### RHO

AACGTGGAGGACTGGTTAGAATCCAATCTTACAGAAAGGAAGTGAACACTATGAGTTTCAAACAGGTGAGACGAAAGGAACTCCGTCTCAAATATGAAAACTTTTTAACCTCTCTTGAGATGAAGTACACCGAGGACAAACTAAATGACCTAGTGTCGAGGCAGAACCAATCACCCGACAGCTCCATTATCTACTGGGTTAACCTTAACTGGCTAACTCGGGAGGAGTTCATTGCTTTAGCCTTTGTAGCCTGGCTAACAGACTCCAAAGCAATTAAGTCCCTCATCCTTGAAGCACTAGTCACTAGGCTAGAAAAGCAAGAAGGTCTGAGTGGAATGGAGAAATGGAATTTAGCGAGATTTCTCTCGGACCACGTTAAACGTGGACTTCCATTTTACCGATCCCACCAGAAAGTCTTCTTTAGCTATCTACCGAAGCGACAGGTTCTTGGATTTCTTGGTGACCCAAAGCTCCAGAGATCCATTAACCGGCGACTCAGACCAAAGGTCTGGACCCCGCCGAAGCCGAAAAGGACTCAGAGAATTAGAGGCTATCGGGATCATGGTACCTGTCGACCGAGTCATAAGTGGACTCCGCGGGACATCATCTCTCCGAGGGAAGCTTTCAATCGAGAAGAGCTCAAACGACTCAACGCTGAATGGCTCATTAGGTGTTGCACTGAACCTTACTGGAATTGGTTCTGGCCAAATCCACACAAACGTTCCAGTGCTTAGTTAACTGCAGGCCATGATCCTTAGGGACTCTCGGGTCTCCTAAGGTGATTGGTTCTGTAAGTTAACTAGCCTCCTTG

#### SIGMA

GCTGTGTTCCTCTCTGACCGAACTTCCAACAGAAAGGAAGAGTAAACACAGTGCTAAGTCTTACTCTCGATGAGTTCGACTTGCAACTGAACAATTTTTACTCTAAGAATCTGAAGGAGAGAACAGAGAGTCTCCGAACCAAAATCCTGGATAGTCCGACCCA GCAACCTGAAGGCCTGGATCTCACGCGGTTAGTGATCAAGGCAAGAAAACAGAGACACTGGATAGGATTAGCTCTTCTGTCTTGGTACATGCCAGAAGAGGTCAGGTTCCTCCTCCAGCTCAAGTTACATGAGCACAAAAACTGGGACGTAGTGGATTTTGATCTTGGAGAAATTCTCTTAAGTTCAAAGTCTCACGCCCTCGCTTTTGTCCTCGAACAAGAGTGGTGGAATGAATGTGACTTTTTCGGGAATGTCCTTGATAGGGAGTTAATCTCAATCTGGCGTCGTGTTGACTTTGTCAAGATCTCTACTCGCAGAGTTCCGAGGTATTCCGGTTACTGCTGTGGCTACCAGGATTCTAGCCGGAGAGCTCCTAGTCCTCTTCCTCTAGAACTTAGAGCAAGAAGTTCAGTAGCAGAAGAACTAGAGAGACAAGAAAGAGCACAGGTTCACCTTCTAATCTGGAAGAAGAAGGTCGAGAAAAGAATCACTGCTTAATCTTTCTCTCGGGAGAACGGGGTCGGGCGCACCGGCCCCCTCTCTTTTGTACCTTATAGAAGAATGTTAACCGCAACTCATTGAAAAGTCTAAGCAATTCTTCCAGTAAGTTAACTACCAAAGAAAGTACCTAACAAAGGAGTAAGACATTTTGGAAAAAGTTCTTGTTGAACTAATTTCAGAAGTCGAACTTCTTGTGAGGACTTCCGGAGTTACTCCTGAAAGTCTTGACCTAGTCTTCTCGCTGAGATTGCTGGTTAACTCCTTCATAAGGACGGAGAGTGTGGGACCAGCGGCCACATCCGATCTTCTGATCAGATTG ACTAG

#### TAU

ACGATTTTTTTTGTAATCTAGTGCACTTTGAATCGGAAGGGCAACGGACTATGAATCTCGAGTTGTCACGCTGGTATGAAAGCGTGGGAAGGACACATTGCGATAGTTTTATCGGTAGACTCGAGGCCCTCAGCACCAAAGTCACGAATAACTCCAAAAAGGAGCCCTGGTTCCTTCACTTTGCCTGGAGAAATGTTAGTGATAGACAAGCGCTCAATCTAGCTCTTATGATGAACTATCTGAACTTACATCCTGAGCTCGTCTTAAGCAGGGATAAGACCTTTCAATTTCTCGCCTTGGAAGCGATAGACTCTATCAAAAGAACACTACGCAGATTCCAGGACATCAAGAAGATGCCCTTGATTATAAGTGTAGTTCTAGTTGATAAAGGTGTATCGTTCACCTATTTGAGTNNNATTGAAGGAGCTTACCTTCCTAAAAGAGAGTTCAAGGGTAACTTTCTAATAGCTAGTCATAGAATTGTAGATGAACTACTTTCTATACGCTATCAAACTAACAGGAGACTGAAGAAGCAGGAGTTCCATAGAGGTTATCGTGACCATGGCTCGATGGCTTCGGTACACGATAGAGCTAGACGCAGTGCGAACACCTCTCACGATATTCTACTAAACGAACAACTAGAAAACATAAGATCTGCACCCGATCCTCTCAAATGGGCCTACGACCACAAAATTATTCCGGACGAAAGATCAACTTAGATAATTCAGACATAAATTCCTCGGACACTCAAGGGTGTTCCGAGGGGTTGTGTTCTGAAGTCTAAAATTTGACTCTCGTNCGATTTTTCTTGTAATCTAGTGCACTTTGATCTGGAGGGGCAATAGACTATGAATCTAGAGTTGTCACGCTGGTATGAAAGCGTGGGTAGGGCACACTGCGATAGCTTTATCGGTAAACTCGAAGCCCTCGCACCAAAGTCACGAATAACTCCAAAAAGGAACCCTGGTTCCTTC

#### UPSILON

CTCCGTCTTGGGACAAGGCTACCTGAGGAAGAAAGAAAGGACTAGTTGAAATGATGAATTTAGGTCTTAATGACCTAGATTCTCAAATTCAGAACTTCCTTTCTAAACTTTCTTTGGGAAAGCCAGAGAGTCTCGAACGGTCACTCTCGGATAGTCCGACACAGTACTCAGCTGTACCGGATTACGACTTCACTCAGATTTTTCTGAAAAGTCGACGGAAGACCGATCTACGAAGACTTGCCATCTTATCTTGGTACTTTCCGGAAAAAGTAAGAATCATCTTACAGTTATCTCTGAAAGAATACTGGGACAAGAGCGACCAGTTTTGTGTAGATAAGGAGATTCTCCTTGTCTCCAAAGCGGTCGCCTTGGCCTGGATTCTTAGTGAGAGTAACTGGAGTGAATCTGACTTTTTCGGAAACATCCTTAATAAGAAGGAA

TGTCAACGTTTGGTCGATTCCCTTGACTTCTTTAGAAATTCTAAGAGAAGGGTAAAGAGGTACACTGGATACTGCCGTGGCTACCGAGAGAGTAACCGCCGAGCTCCTAGGTCTTTTCCTAGAGAGTTGGAAAGGTGGGTTCTGGATGAGGAAATCTGGGATCAAAAAGTACTAAATTATCAAATCAAAGTCCAATCTATTCTCTCCAGAATAGACAAGTTCCTTGAGGAAGTAGTCTAGTACTCTCAAAGGGATCGGTTCTCCGACCCCTTTGAGTTTTGGCGATCTTGTGATCGTTCTAGGAGGTAAAAGAATGAAGGGAATGAATGATGATGGAACAAAAGTTCTACTTGAAGGAATTGAAGACATTGTCTTCTACCTAGAAGGTAGAACAAATTGGACCATCTCAGACATCCTCATCCACTCTTATACCAGCCAGATCGTCACTAAGATCCCGGCTCAAAGTGGGGACGGGAACACCTCCTTGAGGGTCC GACTC

### Oblin-1 Consensus

Oblin-1 protein obtained from the consensus of each cluster:

>ALPHA Oblin-1

MLNFLEDTTLYHSRDLFGNILPTSPEKQVRLANLVSVRYGPTSLPKRVIRRRGYKDHGSRRFPHEVHDLSSGKLAQIKYEEEIQSYHDTLLFLRGWLDGF

>BETA Oblin-1

MLEVRFSTLENQIRNFYNLFLKKECEDRSRSVVDSSNLETDQFEPNGLSFASNLSLTEKILLGVFQWYMPKELGVLTNLWLEEHWGGEFKEIKAVLLNSKDTALGYLLVSDRWSTRDFYGNILKENNVKRILLSMKVRRRNKSRPKRKVWRRGYNDKGSTRLPHDSVRFDYRKLKSVIQTIEEQELLDSKIESLFFIIGQRIEKDEILDYYDSIEF

>CHI Oblin-1

MRDKNLTLKQFNIQLERKIHQGNPFDERFLTRPVFVSPTRPIDPSEVSIRYPRDFRQLRSLCILQWYVPETLHWRMFLDLKDFSFSQLNEKQRIEIQILLSSKEDMISYLYFTERYSGSEIFGNILGNDLTDLKKSMIFKRLKYKRPKRAIRRRGYKDKGSRRPDHRWLPKEDFTFTELQIEKEKRNYQLQTITNRILKTLRQQLKS

>OMEGA Oblin-1

MKLSCDSALPETRNLQKEIVRNLTEQFKDRRLRGLVFRPSLGNFDPEHVKYSKRYLDLVTLLCICYFSDLLDPLDWYLRISVSEYLFKKLDFPELYASTISRRVLLEVISGTISTISPNVFFGNCLRQQNLERVLKNLYLSRNLPVRLKIPVRRRGYKDHGSLRPNERWIENHDYSFSELQFELEQERQDYINLVTSVLNTIQGSSNSNSSIFENLRKECLEWHLKKQEITSTSQK KLRKTNNC

>OMICRON Oblin-1

MRIKLNNLDVLLNTSNLPYRRNITSTERKQIDRILNVYKEAYKYQGTRISEVTFDPNFSERDIFILLSSFCPEVIEFGLQILCFQGELSLNSESDTSILLGILASEPRLLPILLEYKGWRQMPDVSQTLEKIRVQKIYRPRIEQVQRKRGYTDHGSMAPLETKIRKSCLTDELLEEYQRRVKYDRVLKDTIDFLVGYYS

>PHI Oblin-1

MISWFSDDISRCYLQLWLSEQKSLSGDYKVLLDLVLKSKVIALMWFSERYHTREFFGNIKKRMQQIIQMVKPITKIPRKPKRSVWRRGYRDHGSRRPPEKWVETSDWTFSQLQEEIERYRQSLLDTADFLEGWLY

>PI Oblin-1

MKMNRHIVSSQFDLMNVPGVERIKSIALESYQEVDPNKTLISEADLFGTTLPDLVYKGTVLKSIRLRYTKKVKYITLVRFLTGKICFDSERVQRIDLLLIYDSMLILQDLVEKDENFKVKFGSDLESLAKILKSFKLHPKTKILDVKKLGNQIEKEIPNFVLPKRNLSTIWKYVEKMYFFSPSTSSGVELKRLPPKLYIGKGYTDKGTARNPALDGSPPWQEVGQTLAFDPEEPVDET KRATKK

>PSI Oblin-1

MKETEKLPSEETPLLDDYLGLGKTVPVTSLERNNDAEAQIEPHRFAPKSSRRNDHTQMTKTSLLYVLFEYCIANSLLEFRFFDSCSTIWKLDKFANASFHSSFQKSVSEWRTKIIEDEILRKRLLKQHRMELIRLTYPLPKKRGNSPKRKRGYSDKGSTRPTHQRFRTDKDTTVSVYAEDLRVVRKVKIYGRRPTVTYQRLPYSEEIGRLLELGLLRREGDFMVPIRPKED

>RHO Oblin-1

MSFKQVRRKELRLKYENFLTSLEMKYTEDKLNDLVSRQNQSPDSSIIYWVNLNWLTREEFIALAFVAWLTDSKAIKSLILEALVTRLEKQEGLSGMEKWNLARFLSDHVKRGLPFYRSHQKVFFSYLPKRQVLGFLGDPKLQRSINRRLRPKVWTPPKPKRTQRIRGYRDHGTCRPSHKWTPRDIISPREAFNREELKRLNAEWLIRCCTEPYWNWFWPNPHKRSSA

>SIGMA Oblin-1

MPEEVRFLLQLKLHEHKNWDVVDFDLGEILLSSKSHALAFVLEQEWWNECDFFGNVLDRELISIWRRVDFVKISTRRVPRYSGYCCGYQDSSRRAPSPLPLELRARSSVAEELERQERAQVHLLIWKKKVEKRITA

>TAU Oblin-1

MNLELSRWYESVGRTHCDSFIGRLEALSTKVTNNSKKEPWFLHFAWRNVSDRQALNLALMMNYLNLHPELVLSRDKTFQFLALEAIDSIKRTLRRFQDIKKMPLIISVVLVDKGVSFTYLSIEGAYLPKREFKGNFLIASHRIVDELLSIRYQTNRRLKKQEFHRGYRDHGSMASVHDRARRSANTSHDILLNEQLENIRSAPDPLKWAYDHKIIPDERST

>UPSILON Oblin-1

MMNLGLNDLDSQIQNFLSKLSLGKPESLERSLSDSPTQYSAVPDYDFTQIFLKSRRKTDLRRLAILSWYFPEKVRIILQLSLKEYWDKSDQFCVDKEILLVSKAVALAWILSESNWSESDFFGNILNKKECQRLVDSLDFFRNSKRRVKRYTGYCRGYRESNRRAPRSFPRELERWVLDEEIWDQKVLNYQIKVQSILSRIDKFLEEVV

### Oblin-2 Consensus

Oblin-2 protein obtained from the consensus of the alpha cluster:

>ALPHA Oblin-2 MDSVQILRKKIQKNEEQREFLLKKIGNLEYEINNLEHKIENQQRVLQNLLKEK

### Unknown ORF Consensus

Unknown ORFs found in the consensus sequences of each cluster:

>BETA Unknown 1

MTKRETKIQNLQKLVEDIEKKVEKLEAQKKVYLLTIESLKSKLDSKSDDERK

>CHI Unknown 1

MRVDLLIRYLSTSINRLQLVSNVLMKENQKLTARNSNNENLDDTRHDQITGILKEYNGDAINPIDLPVTEIDLEAVRLLTEELETKVRTILLLEENLTKLIEKDK

>OMEGA Unknown 1

MEVYVPMNAGLRIMIILSQNFSSSWNRNGKITSIL

>OMEGA Unknown 2

MAFEETRDYLNISEKVEKNKQLLAKLDQKILNLQKQRSALAAKITNQEGYLNKCGTGKKITKALEILNLKNKVKGIHKSSSGKSKELSR

>OMEGA Unknown 3

MTLDDLFLYFHRLEITGYVLNCAKKGVHRYEVVLRFCSARNKKSAKRNCQKSHRAVQGPQTSRPCIPTLLGKF

>OMICRON Unknown 1

MSTRKLINTKEPEYPKLPLTRTFQNEIFSYYYLPFVQKSSSSVFKFFVFREN

>PHI Unknown 1

MISLVVTYSCGYQNRNHCQEIIKFFWTLSLRAK

>PHI Unknown 2

MAPGVPPRSGWKPQTGPFLNCKKKLRDIDNRYLILLTFLKVGSIDFLATGGEVVLIQP

>PI Unknown 1

MKPNELQKNRRLIFKPEGYTAAELRDCSLADLLFCDMLRELQSHLEPSKFRPKSSNTSKQKESKLTPG

>RHO Unknown 1

MVPVDRVISGLRGTSSLRGKLSIEKSSNDSTLNGSLGVALNLTGIGSGQIHTNVPVLS

>TAU Unknown 1

MKAWEGHIAIVLSVDSRPSAPKSRITPKRSPGSFTLPGEMLVIDKRSI

>TAU Unknown 2

MGLRPQNYSGRKINLDNSDINSSDTQGCSEGLCSEV

### Conserved Region of Oblin-1 Alignment

Conserved region in the alignments made with the Oblin-1 protein consensus sequences from each cluster:

**Figure 1.**
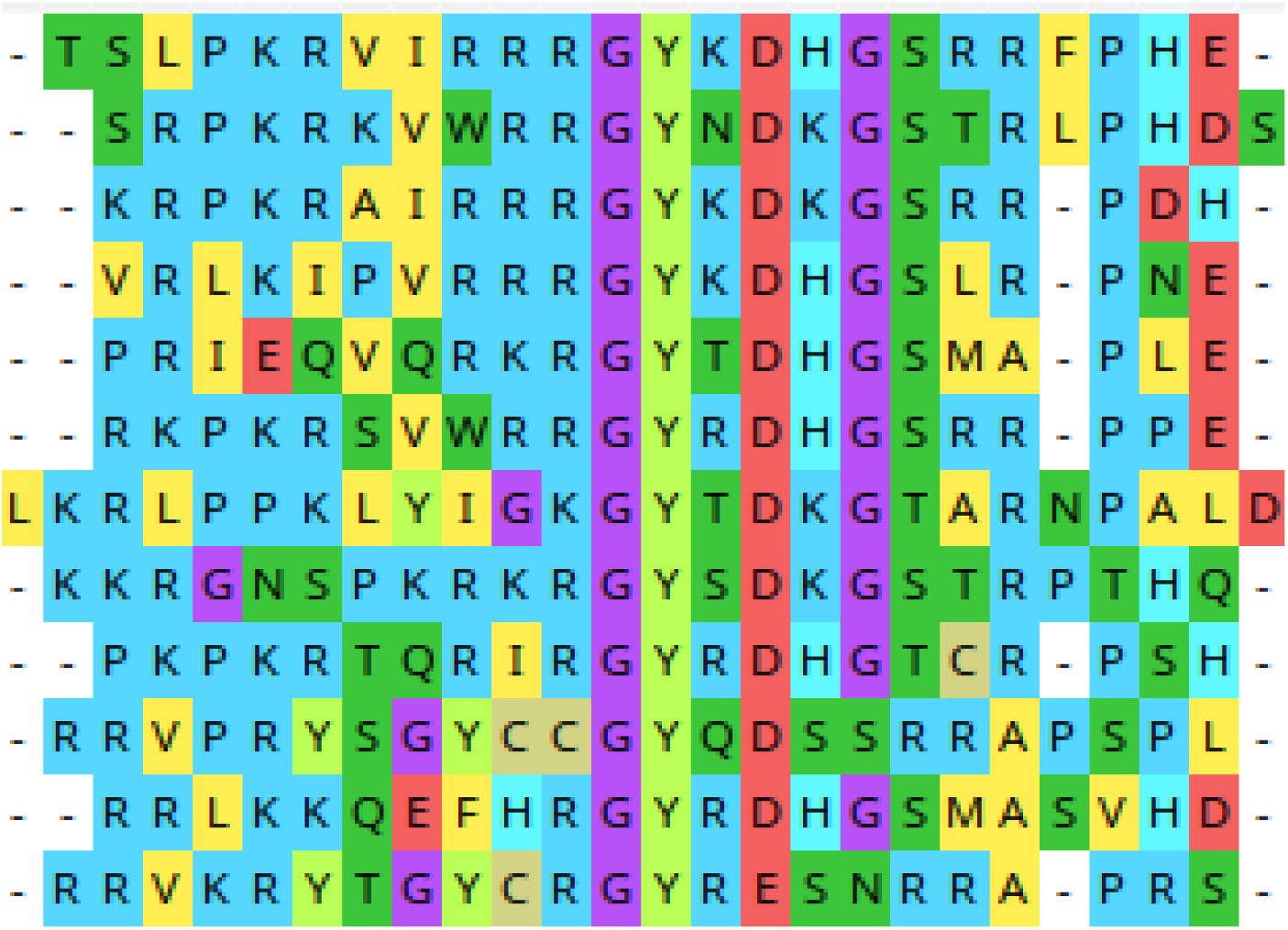
Conserved sequence in the alignment of the consensus sequences from each cluster for the Oblin- 1 protein.

### 3D Structure of the Oblin-2 Consensus from the Alpha Cluster

**Figure 2.**
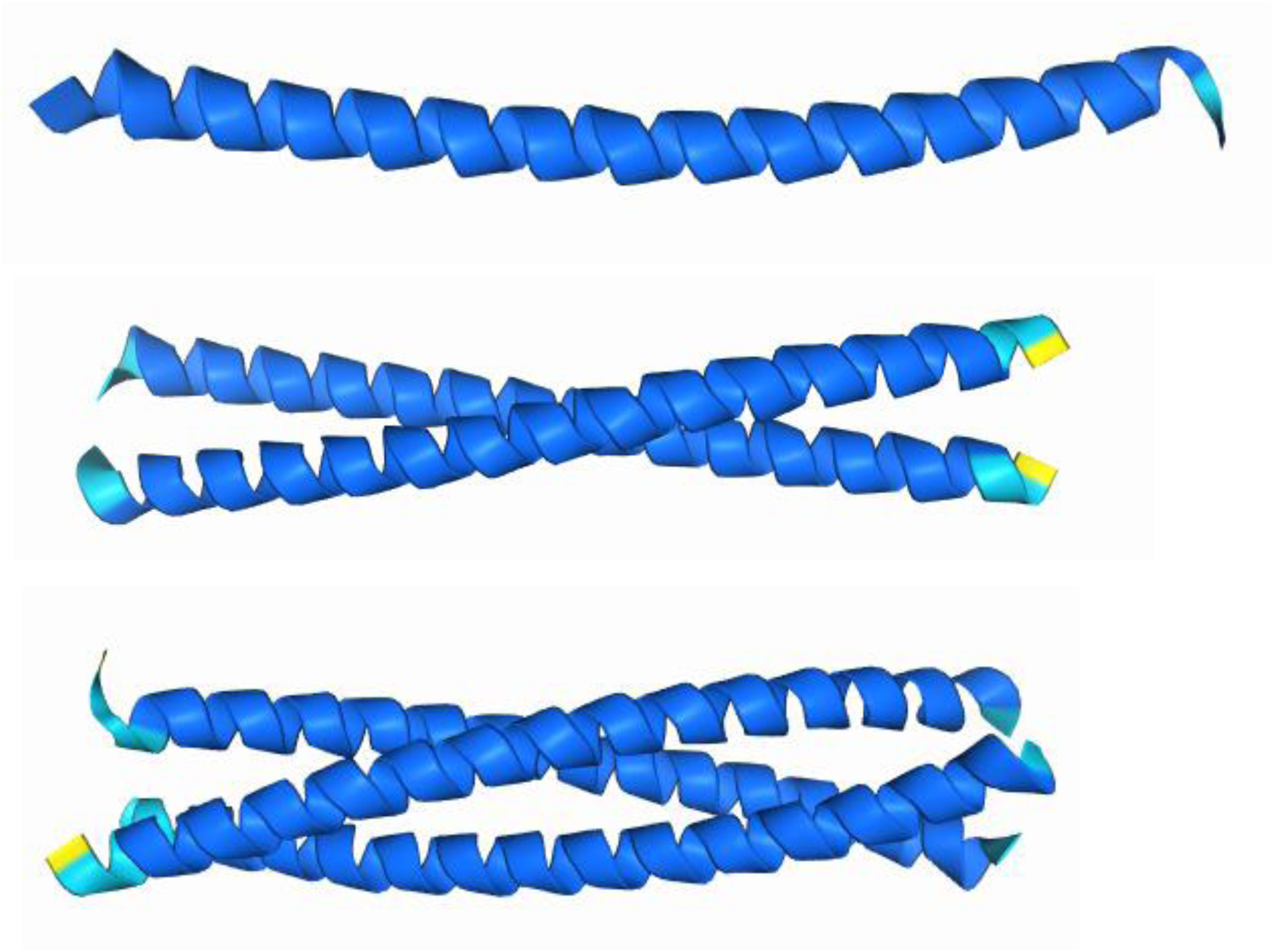
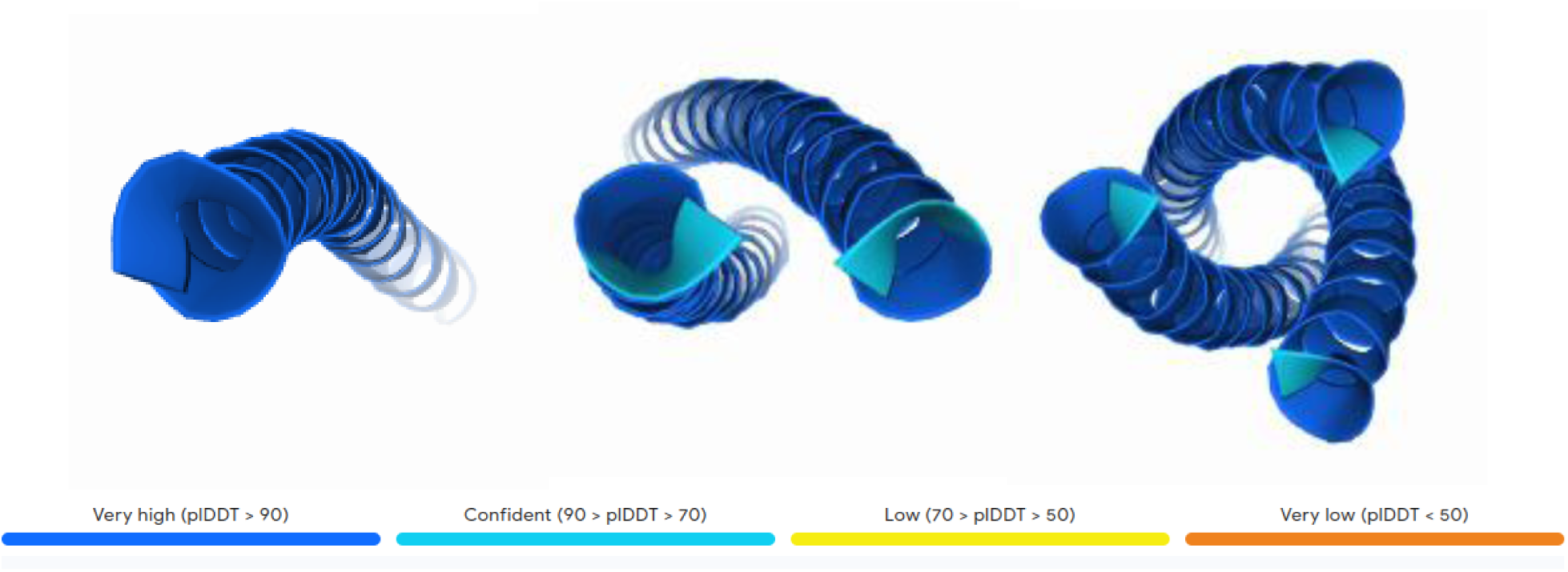
Different 3D structures of the consensus sequence from the alpha cluster for the Oblin-2 protein. Views from two different angles are represented, both as an individual chain and as homodimers and homotrimers. The structure is color-coded from dark blue to orange, indicating decreasing reliability of the 3D structure.

## Discussion

We identified open reading frames (ORFs) corresponding to Oblin-1, Oblin-2, and other unknown ORFs that could potentially be assigned to Oblin-2. Some of these might correspond to small proteins. In line with our findings, for this last concept, Zheludev et al., 2024, established that HHR (hammerhead ribozyme), although not encoding Oblin-2, includes a smaller unrelated ORF.

Through bioinformatics tools, we detected a conserved region among the different consensus sequences of the Oblin-1 clusters, similar to the findings of Zheludev et al., 2024 (RRRGYKDHGSRRFPHEVH). In contrast, the Oblin-1 consensus sequences among different clusters generally exhibit very high variability.

In our findings, we observed a high level of conservation within each cluster. The most variable cluster is “TAU” with approximately 95% conservation, while the most conserved is “BETA” with 100% conservation (Table 2). Since we analyzed complete sequences and did not focus on the Oblin-1 sequences from the alpha obelisk, which are generally poorly conserved according to Zheludev et al., 2024, we did not verify this result.

**Table 2.** Mutation frequency of each cluster.

| Cluster | Mutation frequency | Mutation frequency % | % Conservation | Media Mut. Seq. | Total Seq. |
| --- | --- | --- | --- | --- | --- |
| ALPHA | 0,01243 | 1,243% | 98,757% | 14,460 | 63 |
| BETA | 0,00000 | 0,000000% | 100% | 0,000 | 12 |
| CHI | 0,00673 | 0,673% | 99,327% | 8,268 | 56 |
| OMEGA | 0,00323 | 0,323% | 99,677% | 4,630 | 54 |
| OMICRON | 0,01313 | 1,313% | 98,687% | 11,094 | 138 |
| PHI | 0,00768 | 0,768% | 99,232% | 6,321 | 56 |
| PI | 0,01656 | 1,656% | 98,344% | 19,483 | 116 |
| PSI | 0,00930 | 0,93% | 99,07% | 10,545 | 55 |
| RHO | 0,00389 | 0,389% | 99,611% | 3,157 | 83 |
| SIGMA | 0,00281 | 0,281% | 99,719% | 2,778 | 63 |
| TAU | 0,04445 | 4,445% | 95,555% | 35,468 | 62 |
| UPSILON | 0,01036 | 1,036% | 98,964% | 9,724 | 58 |

Complementing the analyses of Zheludev et al., 2024, regarding mutations found in each cluster, despite the high conservation, some mutations appeared with frequencies around 50%. In this analysis, 50% is the maximum possible frequency since a consensus sequence of these sequences is used as a reference. It is important to highlight that when constructing the consensus for the “TAU” cluster, several positions showed 50% conservation for two nucleotides (Table 2). This led to using “N” as an unknown nucleotide since both had the same frequency of occurrence.

## Conclusions

This study demonstrates that the clusters of the analyzed obelisks exhibit very high conservation within their respective groups. Despite this, some clusters show highly prevalent point mutations. Additionally, the Oblin-1 protein is highly variable among the consensus sequences of each cluster, though a conserved region was identified. Our analyses successfully replicated the three-dimensional structure from the Oblin-2 consensus sequence of the alpha cluster. Finally, we conclude that the “TAU” cluster is the most variable, whereas the “BETA” cluster is the most conserved among those analyzed.

